# Microgravity enhances the viability of midbrain organoids on the International Space Station

**DOI:** 10.64898/2026.05.04.722620

**Authors:** Elisa Zuccoli, Daniela M. Vega Gutiérrez, Aelyn Chong Castro, Lina M. Amaya Mejía, José I. Delgado-Centeno, Miguel A. Olivares Mendez, Carol Martinez, Jens C. Schwamborn

## Abstract

As human spaceflight becomes increasingly relevant, understanding how microgravity affects the human brain is an important but largely unexplored question, particularly in the context of neuronal function and vulnerability to neurodegeneration. Direct investigation of these processes in humans is not feasible, necessitating the use of physiologically relevant *in vitro* model systems. Three-dimensional human brain organoids recapitulate key aspects of brain development and organization and provide an experimentally accessible platform to study neuronal responses under controlled conditions. Here, within the framework of the student competition “Überflieger 2”, we investigated the effects of long-term microgravity on human midbrain organoids cultured for 40 days aboard the International Space Station (ISS).

Midbrain organoids reproduce essential features of dopaminergic neuron development and are widely used to model Parkinson’s disease and related neurodegenerative processes. To enable spaceflight experiments, we developed and implemented an autonomous culture system adapted to the constraints of the ISS environment. During the mission, a hardware malfunction impaired scheduled medium exchange, introducing an additional metabolic stress condition. Despite these limitations, ISS-cultured organoids remained viable and showed robust neurite outgrowth. Molecular and imaging analyses revealed that exposure to microgravity in combination with nutrient limitation induced a coordinated response involving cytoskeletal remodeling, neuronal plasticity, and selective vulnerability of dopaminergic neurons.

These findings demonstrate that human midbrain organoids can maintain key structural and functional properties under prolonged spaceflight-associated stress while activating adaptive response programs. This work highlights the potential of organoid-based systems to investigate neurobiological effects of microgravity and provides a foundation for future studies addressing mechanisms relevant to neurodegenerative disease.

## Introduction

The increasing frequency and duration of human spaceflight have made the effects of microgravity on the human brain an important emerging biomedical question. In particular, it remains unclear how altered physical conditions in space influence neuronal function and vulnerability, including processes relevant to neurodegenerative diseases. At the same time, these conditions also provide a unique opportunity to probe fundamental mechanisms of neural biology under extreme physical environments not accessible on Earth. Three-dimensional (3D) organoid systems have emerged as a powerful tool for modelling human development, disease mechanisms, and drug screening applications (Lancaster & Knoblich, 2014; Clevers, 2016; Rossi et al., 2018; Xu et al., 2026). Human midbrain organoids recapitulate key aspects of dopaminergic neuron development and are widely used to model Parkinson’s disease (PD) and related neurodegenerative disorders (Monzel et al., 2017; Smits et al., 2020). These systems provide experimentally accessible platforms to study early human brain development and neuronal vulnerability under controlled experimental conditions (Lancaster & Knoblich, 2014; Paşca, 2018).

Despite these advances, conventional organoid culture systems remain subject to fundamental physical and physiological constraints inherent to current *in* vitro culture approaches, which influence tissue architecture, growth dynamics, and cellular organization. Brain organoids often exhibit dense cellular packing compared to the developing human brain, and their size is typically limited to approximately 2-3 mm in diameter (Zagare et al., 2021). Limited oxygen and nutrient diffusion, the absence of functional vascularization, and mechanical constraints contribute to heterogeneous growth and structural variability within organoids (Lancaster & Knoblich, 2014; Kelava & Lancaster, 2016). Several bioengineering strategies have been developed to address these limitations, including vascularized organoid models, endothelial co-culture approaches, and perfusion-based culture platforms (Cakir et al., 2019; Pham et al., 2018; Zimmermann et al., 2025). While these approaches aim to mitigate physical limitations, the broader influence of the physical environment on neuronal tissue development remains comparatively underexplored (Mattei et al., 2018; Mozneb et al., 2025).

Furthermore, conventional tissue culture systems cannot fully reproduce the physical environment experienced by developing neural tissue, which grows within fluid-filled spaces rather than static culture conditions (Rehen et al., 2006). Reduced mechanical loading and altered fluid dynamics have emerged as additional parameters influencing stem cell behaviour and tissue organization, particularly under simulated or real microgravity conditions (Mozneb et al., 2025). Previous studies have shown that altered gravitational environments can influence neural progenitor fate decisions and organoid development, including findings from rotary cell culture systems that simulate microgravity (Mattei et al., 2018). Observations from spaceflight studies further suggest that neural organoids cultured under microgravity may exhibit altered morphology and growth behaviour (Marotta et al., 2024; Space Tango, 2019; O’Neill, 2023). Based on these findings, we hypothesise that long-term culture of midbrain organoids under microgravity (30 days) could reduce physical growth constraints and address limitations observed in conventional organoid culture systems.

The physical difference between gravity and microgravity is considered to have a key impact on the entire process of biological evolution (Kamal et al., 2018). The microgravity environment is characterized by greatly reduced gravitational forces compared to Earth gravity, which represents a permanent physical parameter that cannot simply be switched off (Prasad et al., 2020). On Earth, short-term microgravity can be achieved through free-fall approaches such as drop towers or parabolic flights. However, these platforms provide only brief exposure ranging from 20 seconds to 10 minutes (Ferranti et al., 2020), whereas organoid development requires sustained culture over several weeks. The International Space Station (ISS) provides continuous microgravity conditions and therefore enables long-term experiments investigating human tissue development under altered physical forces, which have been shown to influence stem cell proliferation, differentiation, and tissue organization across multiple stem cell and organoid systems (Rampoldi et al., 2022; Mozneb et al., 2025).

To explore this, we established the BRAINS (Biological Research using Artificial Intelligence for Neuroscience in Space) project as part of the Überflieger 2 student program, coordinated by the German Space Agency (DLR) in cooperation with the Luxembourg Space Agency (LSA) and organized by Yuri GmbH. Healthy human midbrain organoids were cultured aboard the ISS for 30 days using custom hardware designed by Yuri GmbH and integrated into Space Tango’s CubeLab™ platform, which meets NASA’s requirements for spaceflight hardware and containment (Melnik et al., 2021; Shaka et al., 2022). Following return to Earth, organoids were analyzed using high-content imaging and transcriptomic profiling to evaluate structural and molecular responses to the spaceflight environment. Despite a hardware malfunction that prevented scheduled media exchanges, ISS-exposed midbrain organoids remained viable and exhibited robust neurite outgrowth. Molecular analysis revealed that spaceflight-induced microgravity and nutrient stress triggered a coordinated response involving neural plasticity, cytoskeletal remodeling, and selective neuronal vulnerability. These findings suggest that human midbrain organoids can activate adaptive programs that preserve key aspects of neuronal structure and function under extreme environmental stress.

## Materials and methods

### Ethics approval

The work with iPSCs has been approved by the Ethics Review Panel (ERP) of the University of Luxembourg and the national Luxembourgish Research Ethics Committee (CNER, Comité National d’Ethique de Recherche) under CNER No. 201901/01 (ivPD) and No. 202406/03 (AdvanceOrg).

### Midbrain organoid culture

A wild-type induced pluripotent stem cell line (iPSC) was obtained from Reinhardt *et al*., 2013 (Supplementary Table 1). One clone of the iPSC line was used, which was derived into neuroepithelial stem cells (NESCs) as described in Reinhardt *et al*. (2013). NESCs were cultured in N2B27 maintenance media in 6-well Geltrex (Thermo Fisher Scientific, Cat. No. A1413302) precoated plates. The N2B27 base media consists of a 1:1 ratio of DMEM-F12 (Thermo Fisher Scientific, Cat. No. 21331046) and Neurobasal (Thermo Fisher Scientific, Cat. No. 10888022) and is supplemented with 1% GlutaMAX (Thermo Fisher Scientific, Cat. No. 35050061), 1% Penicillin/Streptomycin (Thermo Fisher Scientific, Cat. No. 15140122), 1% B27 supplement w/o Vitamin A (Life Technologies, Cat. No. 12587001) and 2% N2 supplement (Thermo Fisher Scientific, cat. no. 17502001). For the maintenance of NESCs, the N2B27 base media was supplemented with 150µM ascorbic acid (Sigma-Aldrich, Cat. No. A4544), 3µM CHIR-99021 (Axon Medchem, Cat. No. CT 99021), and 0.75µM purmorphamine (Enzo Life Science, Cat. No. ALX-420-045). The NESC maintenance media was exchanged every second day.

Midbrain organoids (MOs) were generated according to Zagare *et al*. (2021). Briefly, NESCs were detached at 80% confluency using Accutase (Sigma-Aldrich, Cat. No. A6964). Cell viability was determined using Trypan Blue, and 9000 live cells were seeded into each well of a 96-well ultra-low attachment plate (faCellitate, Cat. No. F202003) in 150µl of NESC maintenance medium, initiating day 0 of organoid culture. On the 2^nd^ day, media was changed to midbrain patterning medium, supplemented with 200µM ascorbic acid (Sigma-Aldrich, Cat. No. A4544-100G), 500µM dbcAMP (STEMCELL Technologies, Cat. No. 100-0244), 10ng/ml hBDNF (Peprotech, Cat. No. 450-02-1mg), 10ng/ml hGDNF (Peprotech, Cat. No. 450-10-1mg), 1ng/ml TGF-β3 (Peprotech, Cat. No. 100-36E) and 1µM purmorphamine (Enzo Life Science, Cat. No. ALX-420-045). A second patterning medium change was performed at day 5.

On day 8, MOs were embedded in Geltrex and transferred to 24 well ultra-low attachment plates (Celltreat, Cat. No. 229524) on an orbital shaker (80 rpm) at 37°C, 5% CO_2_. From this point, the culture medium was switched to differentiation medium, identical to patterning medium but without purmorphamine. Media was changed every 3 - 4 days thereafter. NESC and midbrain organoid cultures were regularly (once per month) tested for mycoplasma contamination using LookOut® Mycoplasma PCR Detection Kit (Sigma-Aldrich, Cat. No. MP0035-1KT).

### Protocol for Spaceflight Experiment

The batch of human midbrain organoids was generated and processed according to a timeline aligned with the anticipated launch date (Supplementary Table 2). All preparatory steps were performed under sterile conditions and according to the payload integration requirements of NASA and Space Tango.

The work was performed in the NASA Space Station Processing Facility (SSPF) BSL-2 laboratory, ensuring controlled incubator conditions (37°C, 5% CO_2_). Due to a shipment issue from Luxembourg, a small subset of small molecules had to be sourced locally by NASA personnel. However, identical compounds were obtained and used at the same concentration to ensure full compatibility with the original protocol.

MOs were generated as described previously on day 0 and followed the standard patterning schedule until day 6. The final embedding in Geltrex and transfer into Yuri Type-IV devices (yuri003 and yuri004) occurred on day 8. Ten MOs were placed in each device, which was then sealed and transferred for final integration into the CubeLab. 48h later, MOs were launched to the ISS aboard the SpaceX Falcon 9 CRS-27. Lastly, 72 hours after launch, the experiment was installed in the ISS, kept there for 20 days.

### Protocol for Ground Control Experiment

Ground control experiments took place at the Developmental and Cell Biology Lab of LCSB using the same Yuri Type-IV bioreactor hardware and experimental timeline as the spaceflight experiment. MOs were generated as described previously on day 0 and followed the standard patterning schedule until day 6. On day 8, MOs were embedded in Geltrex and transferred into Yuri Type-IV devices (yuri003 and yuri004). Ten MOs were placed into each device. The devices were incubated under standard laboratory conditions (37°C, 5% CO_2_) for the full experimental duration. To ensure comparability with the ISS experiment, ground controls replicated all key operational parameters, including the reduced medium exchange frequency, identical aged medium formulation, and the same autonomous hardware operation sequence. In addition, post-experiment handling, including delayed fixation and transport-like storage conditions, was reproduced to match the spaceflight return timeline. Routine mycoplasma testing was performed as described for all organoid cultures.

### LDH Assay

LDH activity was measured using the Promega LDH-Glo™ Cytotoxicity Assay Kit following the manufacturer protocol (Promega, Cat. No. J2381). Briefly, 25μl cell culture supernatant was collected from each sample and mixed with 25μl LDH Detection Reagent per well. The reagent was prepared previously by mixing the Reductase Substrate 1:200 with the LDH Detection Enzyme Mix. The sample-reagent mix was incubated for 60 minutes at RT and luminescence was measured with the BioTek Cytation 5 Cell Imaging Multi-Mode Reader (RRID:SCR_019732). Cytotoxicity percentage was calculated following the manufacturer protocol.

### Immunofluorescence staining of midbrain organoid sections

Midbrain organoids were fixed with 4% PFA overnight at 4°C and washed in 1x PBS. They were then embedded in 3 % low-melting point agarose (Biozym Scientific GmbH, Cat. No. 840100) in 1x PBS and sectioned into 60µM thick slices using the Vibratome VT1000S Leica Microsystems (RRID:SCR_016495). The sections were permeabilized for 30min in 0.5% Triton X-100 at RT, followed by one quick wash with 0.01% Triton X-100 in PBS. Sections were then blocked for 2h at RT with blocking buffer containing 2.5% BSA, 2.5% donkey serum, 0.01% Triton X-100 and 0.1% sodium azide in PBS. The antibodies TH (rabbit, Abcam, Cat. No. ab112, RRID:AB_297840) and MAP2 (chicken, Abcam, Cat. No. ab92434, RRID:AB_2138147) were diluted in blocking buffer and the sections were incubated with the primary antibody dilution for 48h at 4°C on an orbital shaker. After the incubation, sections were washed three times for 5 minutes with 0.01% Triton X-100. Subsequently, they were incubated with the Alexa Fluor^®^ conjugated secondary antibodies donkey anti-rabbit 568 (1:1000, Thermo Fisher Scientific, Cat. No. A10042, RRID:AB_2534017), donkey anti-chicken 647 (1:1000, Thermo Fisher Scientific, Cat. No. A78952, RRID:AB_2921074), and the nuclei stain Hoechst 33342 (1:1000, Invitrogen, Cat. No. 6224) for 2 hours at RT on an orbital shaker. The washing step was repeated, and sections were washed once with MilliQ water. They were mounted on slides (De Beer Medicals, Cat. No. BM-9244) and covered by Fluoromount-G® (SouthernBiotech, Cat. No. 0100-01) mounting media.

The TUNEL In Situ Cell Death Detection Kit, Fluorescein 488 (Merck, Cat. No. 11684795910) was used to stain apoptotic cells. The components of the kit were incorporated into the staining procedure, with the secondary antibody diluted in the buffer provided by the TUNEL assay kit. The sections were incubated for one hour at RT on an orbital shaker.

### Image acquisition and analysis

High-content imaging was performed using the Yokogawa CellVoyager CV8000 (RRID:SCR_023270) equipped with a 20x objective. One to three sections from at least three organoids were analysed per condition and staining combination. The acquired images were processed with a customized analysis pipeline in MATLAB (2021a, Mathworks, RRID:SCR_001622), following the methodologies outlined in Bolognin *et al*. (2019) and Monzel et al. (2020). Sections exhibiting suboptimal masking for target proteins or those that were out of focus were excluded from further analysis. Data normalization was performed by calculating the average value per feature and per time point across batches. Outliers were identified and removed using the interquartile range (IQR) method, based on the 25th and 75th percentiles. Qualitative images were captured using a Zeiss LSM 710 Confocal Inverted Microscope (RRID:SCR_018063) with 20x, or 60x objectives, and processed using the ZEISS ZEM Microscopy Software (RRID:SCR_013672).

### RNA extraction, library preparation and sequencing

Total RNA was extracted from two to three organoids per condition using the RNeasy Mini Kit (Qiagen, Cat. No. 74106) according to the manufacturer’s instructions. Messenger RNA (mRNA) was isolated using poly-T oligo-attached magnetic beads and subsequently fragmented. First-strand cDNA synthesis was performed with random hexamer primers, followed by second-strand synthesis using either dUTP (for directional libraries) or dTTP (for non-directional libraries). Libraries were prepared and sequenced on an Illumina platform by Novogene.

### Transcriptomic analysis

RNA sequencing data were processed using Galaxy server v23.2.rc1 (RRID:SCR_006281), following established Galaxy training tutorials (Hiltemann et al., 2023; Doyle *et al*., 2024). Reads were aligned to the human reference genome (hg38) using HISAT2 (RRID:SCR_015530). Aligned reads were quantified with featureCounts (RRID:SCR_012919), and differential expression analysis was conducted in R Project for Statistical Computing (v4.4.2, RRID:SCR_001905) using the DESeq2 package (v1.42.1, RRID:SCR_015687) (Love et al., 2014). P-values were adjusted for multiple testing using the Benjamini– Hochberg method (Benjamini & Hochberg, 2005). Pathway enrichment analysis was performed using EnrichR (RRID:SCR_001575) (Kuleshov et al., 2016) on DESeq2 output, with reference to the Reactome Pathways 2024 (RRID:SCR_003485) (Milacic et al., 2024), to the KEGG 2021 Human Pathways (RRID:SCR_012773) (Kanehisa & Goto, 2000) and Gene Ontology (GO) Biological Process 2023 (RRID:SCR_002811) (Ashburner et al., 2000; Aleksander et al., 2023) gene set libraries.

### Data processing and statistical analysis

Data was analysed and visualized in GraphPad Prism (v.10.2.3, RRID:SCR_002798) or R statistical programming software v4.4.2. (RRID:SCR_001905). Statistical significance was assessed according to the conditions to be analysed. An ordinary two-way ANOVA with Šidák’s multiple comparisons test or with Dunnett’s multiple comparison test, as well as a Kruskal-Wallis with Dunn’s multiple comparison test was done in GraphPad. Mann-Whitney *U* test of two conditions was done in GraphPad and in R. Significance asterisks represent * p<0.05, **p<0.01, ***p <0.001, ****p<0.0001, ns stands for not significant. Error bars represent mean + SD.

## Results

### Experimental setup with the Yuri Type-IV bioreactor systems

For this study, we first performed a series of ground-based optimization experiments to determine the environmental and hardware parameters most critical for long-duration culture of human midbrain organoids under autonomous conditions. These tests guided the final configuration of the Yuri Type-IV bioreactor systems (Yuri Gravity, 2023), which were subsequently used both on ground and in space. Each device consisted of an inner shell (Figure 1a) containing two 11⍰mL⍰±⍰0.3mL tanks and a central culture chamber of 13⍰mL⍰±⍰0.3mL, where the 10 organoids were maintained. One tank contained the fixation solution, while the second was divided by a membrane to separate fresh nutrient medium from waste. The inner shell was enclosed by an outer shell (Figure 1b), providing a first layer of biological containment. The complete setup comprised a biological unit (green), a technical unit (blue) containing electronics, and the external CubeLab™ housing (red), which served as a second containment layer and integrates the overall system (Figure 1c). Figure 1d highlights the thermal architecture of the system. A 5⍰mm insulation layer surrounds the bio inner structure, and 5⍰mm extrusions around the screw positions prevent compression of the insulation during assembly. The heating system consisted of two RS Pro 25×50mm silicone heater mats positioned between the inner and outer shells to ensure uniform thermal distribution. The full Type-IV unit weighed 723.65g, increasing to 1,820.87g when integrated into the CubeLab™.

**Figure 1.**
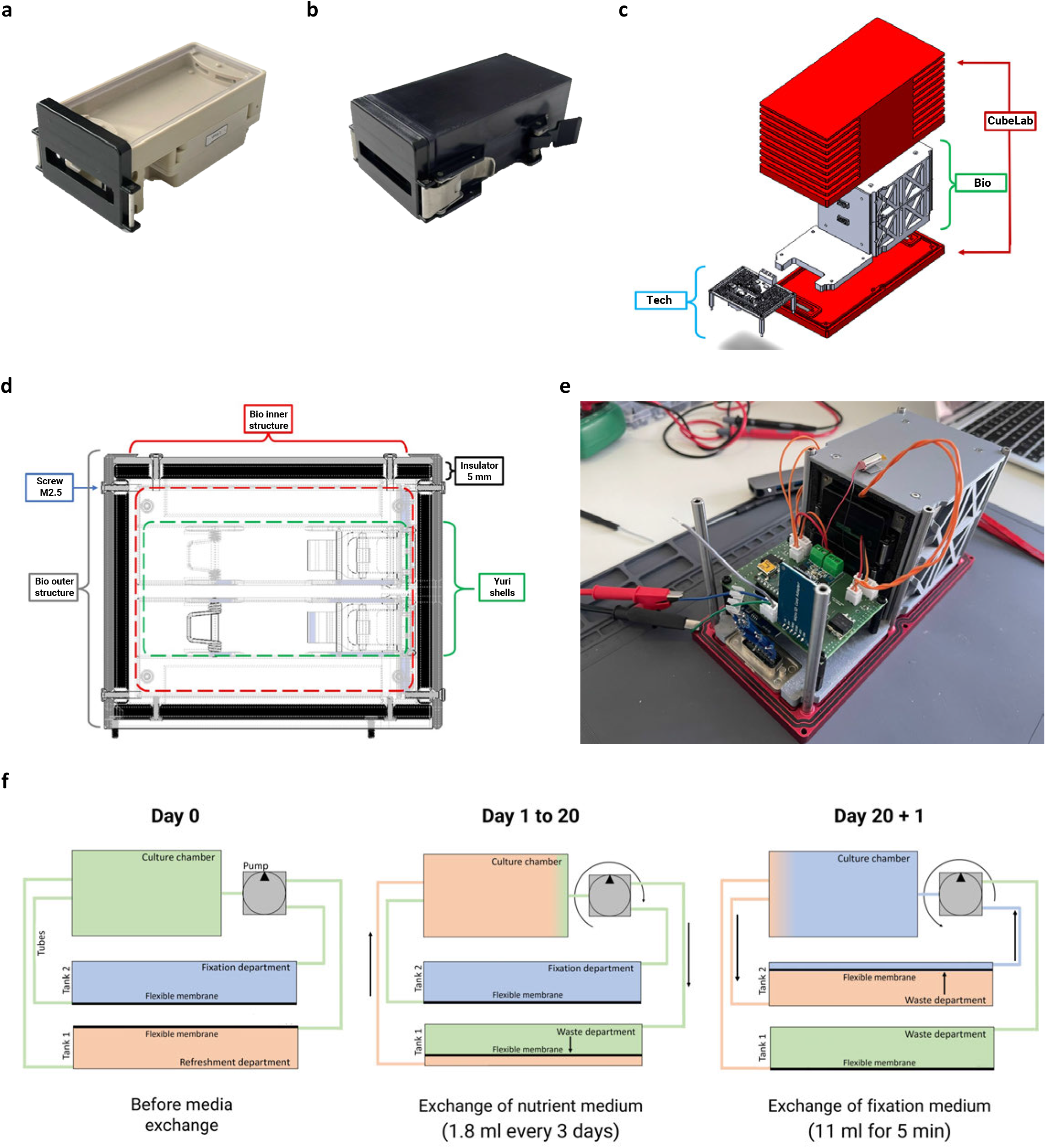
Experimental setup with the Yuri Type-IV bioreactor systems. (a) Yuri Type-IV inner shell and (b) outer shell. (c) Exploded view of the assembly with the technical (blue) and biological (green) components inside the CubeLab™ (red), (d) Thermal insulation (5mm thickness) surrounding the bio-inner structure (red) with the shells (green), the M2.5 screws (blue) surrounded by the bio-outer structure (grey), and (e) the finished device before launch. (f) Media pump and extraction schematic. (f) Software flow chart with commands: start/ed actions (red), nutrient exchange (green), temperature control (yellow), fixation (blue) and data storage (purple).

Fluid handling was controlled by an autonomous microcontroller (Arduino Nano) programmed to perform partial media exchange every 3 days by delivering approximately 1.8mL of fresh medium into the culture chamber while removing an equivalent volume of waste. A multi-timer system coordinated all onboard processes. Temperature was set to be recorded every second and logged to the micro-SD card every minute, fluidic exchange was scheduled to occur every 72⍰hours, and motor activation durations were calibrated to ~ 27⍰seconds per 1mL delivered. The operational logic of the media-exchange and waste-removal system is illustrated in Figure 1f, which shows the pumping and extraction cycles that govern nutrient renewal throughout the mission. The schematic shows an automated pump cycle in which waste is first removed before fresh medium is dispensed, with each motor timed according to its experimentally determined flow rate, ensuring precise volumetric exchange under autonomous conditions. On the final day, the chamber cools from 37°C to ~ 25°C, triggering an automated 11mL fixation perfusion (~ 5min) after which all motors and heaters deactivate.

Before conducting the temperature-control performance test, we first validated the basic functionality of the key electronic subsystems required for autonomous device operation. Using simple Arduino-based test circuits (Supplementary Figure 1a-c), we confirmed correct operation of the temperature sensors (Test 1, Supplementary Figure 1a), heater-switching circuit (Test 2, Supplementary Figure 1b), and bidirectional motor control (Test 3, Supplementary Figure 1c). In addition, the real-time clock and SD-card modules were tested to ensure that scheduled actions and temperature logging operated reliably during short ground-based runs. A subsequent breadboard integration test further demonstrated that temperature readout, heater activation, pump commands, and data logging could run together in the intended sequence.

To evaluate the thermal performance and timing accuracy of the finalized hardware configuration, we conducted a full hardware performance test using the complete setup, including both shells with attached heater mats, the integrated insulation layer, and the assembled electronics subsystem (Figure 2a). The electronics consisted of custom PCBs with soldered connectors and an external 12 V power supply, while all 5V components, including the Arduino microcontroller, were powered by the internal regulation circuit. Temperature profiles from both shells showed nearly identical behaviour, reaching the target 37°C setpoint within approximately 39 minutes and maintaining stable conditions with minimal fluctuation (Figure 2b). These results confirmed that the temperature control system performed as expected for a 37°C setpoint and could maintain the necessary thermal stability to execute the planned biological operations. After heater deactivation, the system cooled to ~ 23°C whin 1 hour and 32 minutes, demonstrating sufficient cooling efficiency to support automated fixation protocol. Although the insulation efficiency minimized heat loss during warm phase, the cooling rate remained fast enough to reach fixation conditions within the programmed time window. Quantitative performance metrics, including maximum and minimum controlled temperatures at both 37°C and ambient setpoints, matched estimated values closely (Figure 2b). The corresponding software flowchart (Figure 2c) outlines the control logic underlying these operations: continuous temperature monitoring and data logging, timed media-exchange events, heater deactivation on Day 30, and the initiation of the fixation routine once the chamber reaches the required temperature threshold. Together these results confirm that the thermal control system, fluidic timing, and software logic perform reliably under operation conditions on ground, ensuring robust autonomous function during spaceflight.

**Figure 2.**
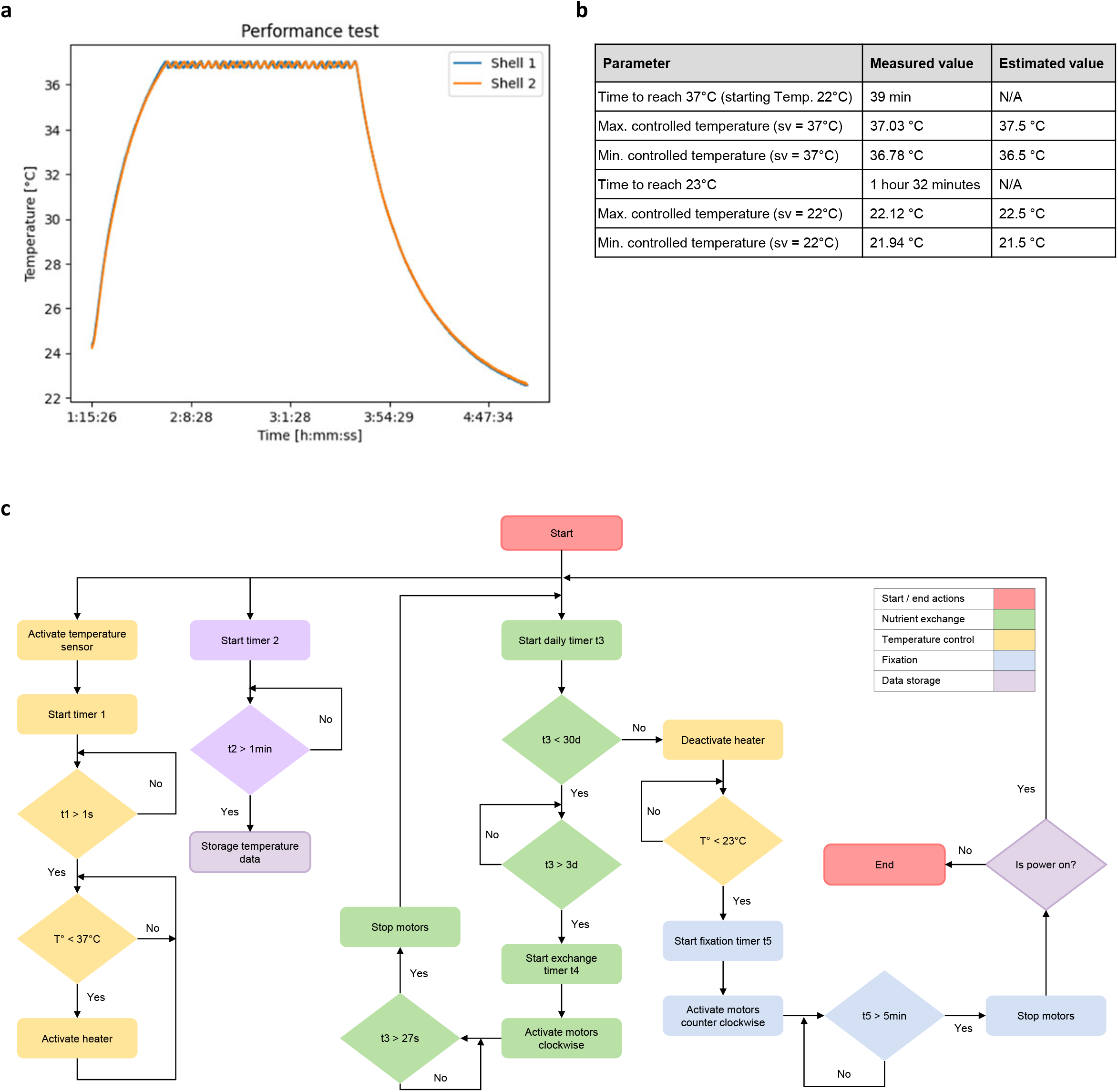
Thermal performance and control logic of the hardware. (a) Temperature profiles of both shells during the ground-based performance test, showing heating to 37°C within ~ 39min and stable temperature maintenance. (b) Summary table of measured versus estimated thermal-control parameters, including heating and cooling times and controlled temperature ranges. (c) Software flow chart outlining the autonomous control logic, including temperature regulation (yellow), scheduled media-exchange events (green), fixation sequence (blue), and data-logging routines (purple).

### Viability and differentiation of Midbrain organoids under simulated spaceflight culture conditions

To assess how medium stability, reduced exchange volume, and restricted exchange frequency may affect organoid viability and neuronal differentiation under spaceflight-like constraints, we performed an on-ground pre-test using scaled-down partial medium exchanges. A volume of 160μL was used to reflect, in 24-well format, the relative exchange volume (~1.8mL) delivered by the Type-IV device on orbit. Medium exchanges were performed either once per week (1x per week) or twice per week (2x per week). Two medium conditions were compared, consisting of a freshly supplemented control medium stored at 4°C and a pre-supplemented aged medium stored at 37 °C to mimic the prolonged warm-storage conditions expected inside the CubeLab™ during the ISS mission. Full medium exchanges (2mL, 2x per week) served as the standard control (CTRL). Organoids were fixed on days 10, 20 and 30 and assessed by confocal imaging of dopaminergic neurons (TH) and mature neurons (MAP2) (Supplementary Figure 2a–c). Cytotoxicity was quantified by LDH release from snap-frozen supernatants (Supplementary Figure 2d).

At day 10, organoids cultured in control medium behaved similarly under both full exchange (CTRL) and partial exchange twice per week (Supplementary Figure 2a, left panel). Since these two conditions show similar morphology and TH/MAP2 staining, the CTRL group was not included in day 20 and day 30 imaging panels. In contrast, organoids cultured in aged medium with only one partial exchange per week (1x per week) showed severe structural loss and almost complete disappearance of MAP2 and TH signals already by day 10 (Supplementary Figure 2a, right panel). This condition was therefore discontinued and not imaged at later time points. Only the aged-medium condition with two partial exchanges per week (2x per week) preserved sufficient structure and neuronal marker expression to follow over time. Organoids cultured in this condition maintained overall integrity through day 20, with clear deterioration by day 30 (Supplementary Figure 2b-c). These imaging observations were consistent with LDH measurements, which showed significantly higher cytotoxicity in aged medium, especially under low exchange frequency (Supplementary Figure 2d).

Although viability under aged-medium conditions did not match the laboratory control (CTRL), aged medium with partial exchange twice per week represented the most biologically sustainable condition compatible with the hardware constraints (limited medium storage, warm-storage environment, and restricted exchange frequency). This combination therefore served as the operational standard for the ISS experiment.

### Fixation optimization for long-term storage under simulated spaceflight constraints

In a second on-ground pre-test, we evaluated fixation parameters for midbrain organoids to identify conditions compatible with long-duration autonomous culture and delayed sample recovery after spaceflight. We assessed the effects of paraformaldehyde (PFA) concentration, fixation duration, and storage temperature. Midbrain organoids were fixed either on day 20 or day 30 and incubated in 1%, 2%, or 4% PFA for 40 or 50 days at 4°C (Supplementary Figure 3a-b). Across all conditions, TH and MAP2 staining remained detectable, with no evidence of over-fixation even at 4% PFA and the longest incubation time (Supplementary Figure 3b, right panel). However, lower PFA concentrations (1% and 2%) resulted in reduced signal intensity and poorer morphological preservation, indicating insufficient cross-linking for extended storage.

To simulate temperature fluctuations expected during payload return, we also compared storage of fixation medium at 4°C to room temperature (RT) prior to fixation (Supplementary Figure 3c-d). Organoids fixed overnight at either temperature maintained staining integrity, although storage at 4°C yielded slightly better preservation of structural detail. Based on these results, 4% PFA was selected as the optimal fixation concentration for subsequent space-compatible experiments.

After independently defining culture conditions (aged medium with partial exchanges twice per week) and fixation parameters (4% PFA), we combined these conditions to validate organoid development under mission-relevant constraints. MOs were maintained in Yuri test-devices for 30 days and subsequently fixed and analysed. As shown in Figure 3a, organoids from both test-devices displayed morphology and TH/MAP2 expression comparable to the control cultures (CTRL). These results confirmed that the automated system preserved viability and structural integrity effectively under hardware-constrained conditions.

**Figure 3.**
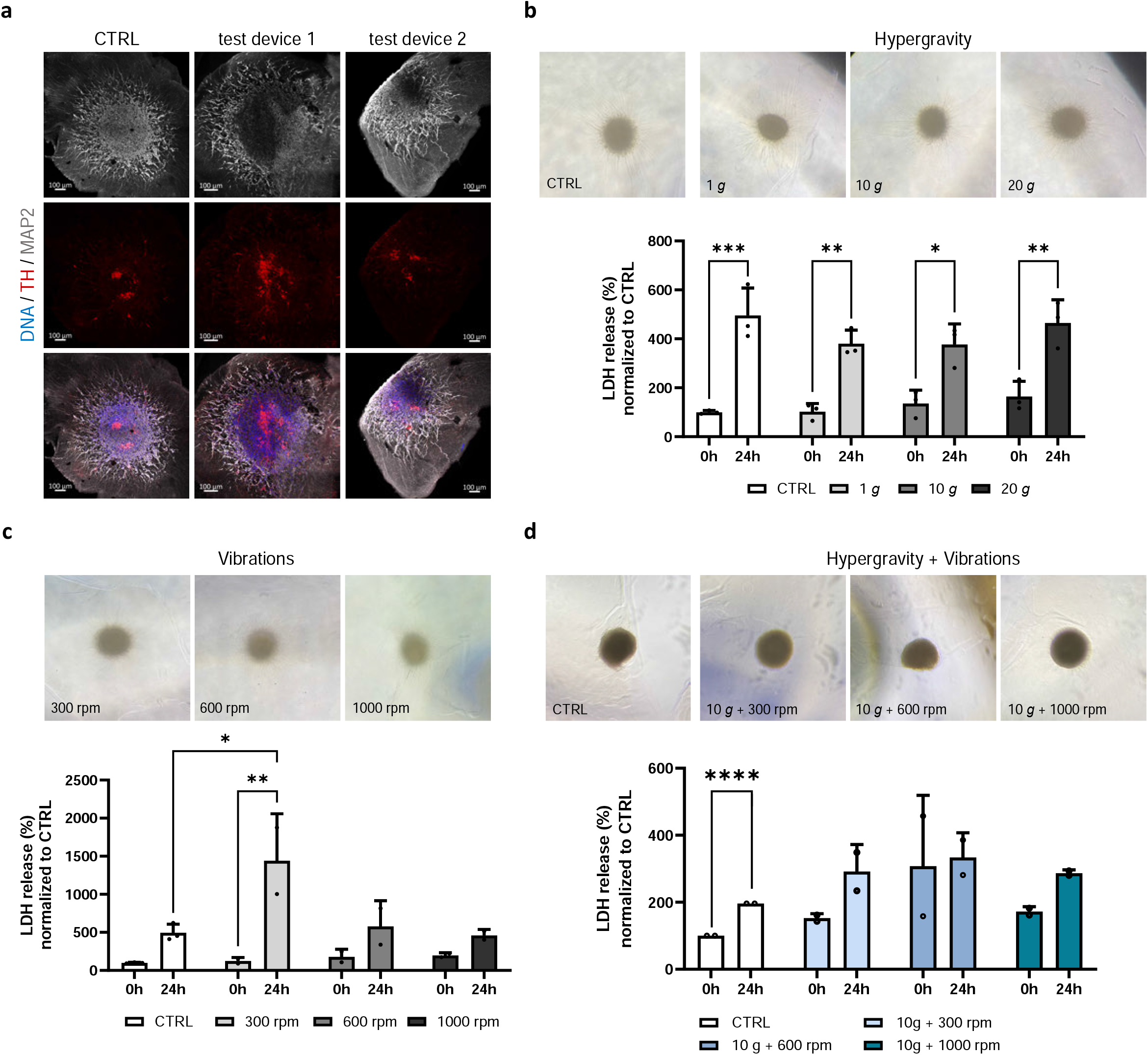
Impact of hypergravity and mechanical vibration on MO viability. (a) Representative immunofluorescence images of MOs cultured for 30 days in the test device. Organoid sections were stained for TH (tyrosine hydroxylase), MAP2 (neuronal marker), and nuclei (Hoechst 33342). Scale bars: 20x, 100μm. (b) Representative brightfield images of CTRL and hypergravity treated MOs. Hypergravity treatment included 1, 10 and 20*g*. Quantification of cytotoxicity by LDH release assay (% normalized to control) for hypergravity in MO media after 0h and 24h. Statistical significance was assessed using an ordinary two-way ANOVA with Šidák’s multiple comparisons test. Data are mean ± SD; *p<0.05, **p < 0.01, ***p < 0.001. (c) Representative brightfield images of CTRL and vibration treated MOs. Vibration treatment included 300, 600 and 1000rpm. Quantification of cytotoxicity by LDH release assay (% normalized to control) for vibration in MO media after 0h and 24h. Statistical significance was assessed using an ordinary two-way ANOVA with Šidák’s multiple comparisons test. Data are mean ± SD; *p<0.05, **p < 0.01. (d) Representative brightfield images of CTRL and hypergravity-vibration combined treated MOs. Treatments included 10 *g* combined with 300, 600 or 1000rpm. Quantification of cytotoxicity by LDH release assay (% normalized to control) for hypergravity-vibration combined treatment in MO media after 0h and 24h. Statistical significance was assessed using an ordinary two-way ANOVA with Šidák’s multiple comparisons test. Data are mean ± SD; ****p<0.0001.

### Effects of hypergravity and vibration on Midbrain organoids viability

To evaluate the resilience of human midbrain organoids to mechanical stressors expected during spaceflight, we performed on-ground tests assessing the effects of hypergravity and vibration using LDH release measured at 0h and 24h post-exposure (Figure 3b–d). Hypergravity conditions were tested at 1*g*, 10*g*, and 20*g* using a centrifuge, representing baseline Earth gravity (1*g*), the upper limit of transient accelerations experienced during rocket launch (~10*g*), and a higher qualification margin (20*g*) typically applied to ensure biological payload robustness (de Sousa *et al*., 2020). Across all conditions, MOs remained morphologically intact (Figure 3b-d, upper panels). LDH levels increased from 0h to 24h in all groups, including the untreated control, reflecting a baseline level of time-dependent spontaneous cytotoxicity during the 24-hour incubation period rather than an effect of mechanical stimulation.

Exposure to hypergravity at 1*g*, 10*g*, or 20*g* did not induce additional LDH release compared with controls (Figure 3b, lower panel), indicating that acceleration forces within this range do not exacerbate cell damage.

To model launch-associated vibration, MOs were exposed to 300, 600, and 1000rpm (Figure 3c). Vibrational testing followed low-, mid-, and high-frequency regimes specified by NASA’s payload vibroacoustic standard (NASA, 2017), capturing representative launch vibration stresses. Among the tested conditions, only exposure to 300 rpm resulted in a significant increase in LDH release at 24h. This effect likely reflects the greater fluid displacement and shear forces at lower shaking frequencies, whereas higher-frequency rotations (600-1000rpm) generate smaller-amplitude oscillations that impose less mechanical strain to the organoids.

We next tested the combined effect of hypergravity (10*g*) and vibration (300-1000rpm) (Figure 3d). Organoids preserved structural integrity under all combined stress conditions (Figure 3d, upper panel). A significant increase in LDH release between 0h and 24h was observed only in the CTRL group, consistent with baseline time-dependent cytotoxicity. No statistically significant increase in LDH was detected in any of the combined stress conditions, though a trend toward higher LDH levels was observed in the 10g⍰+⍰600rpm group (Figure 3d, lower panel).

Together, these results indicate that human midbrain organoid tolerate both hypergravity and vibration across a range of conditions that meet or exceed those during launch. These mechanical stresses are therefore unlikely to compromise organoid viability and structural integrity during CubeLab™ deployment to the ISS.

### Growth and oxygen monitoring of Midbrain organoids in the device

Measuring oxygen consumption is critical to assess the metabolic activity and viability of human midbrain organoids during long-term culture, as oxygen availability directly influences their growth, function, and response to the autonomous conditions within the devices. To assess the feasibility of prolonged culture and monitor oxygen dynamics, we conducted two 20–25-day experiments using the Yuri Type-IV device (Figure 4a). Organoid media exchange was performed twice per week, and oxygen levels were continuously tracked by integrated sensors. In the first run, device malfunctions, leaks, and bubbles caused by improper assembly led to oxygen depletion and likely premature fixative release, halting the experiment on day 10 without immunostaining (Figure 4b). The second run showed improved but still imperfect performance, with medium delivery issues causing oxygen fluctuations that only stabilized after day 14 (Figure 4c). Immunostaining confirmed organoid viability, though some fixation artifacts were noted, possibly due to early washing (Figure 4d).

**Figure 4.**
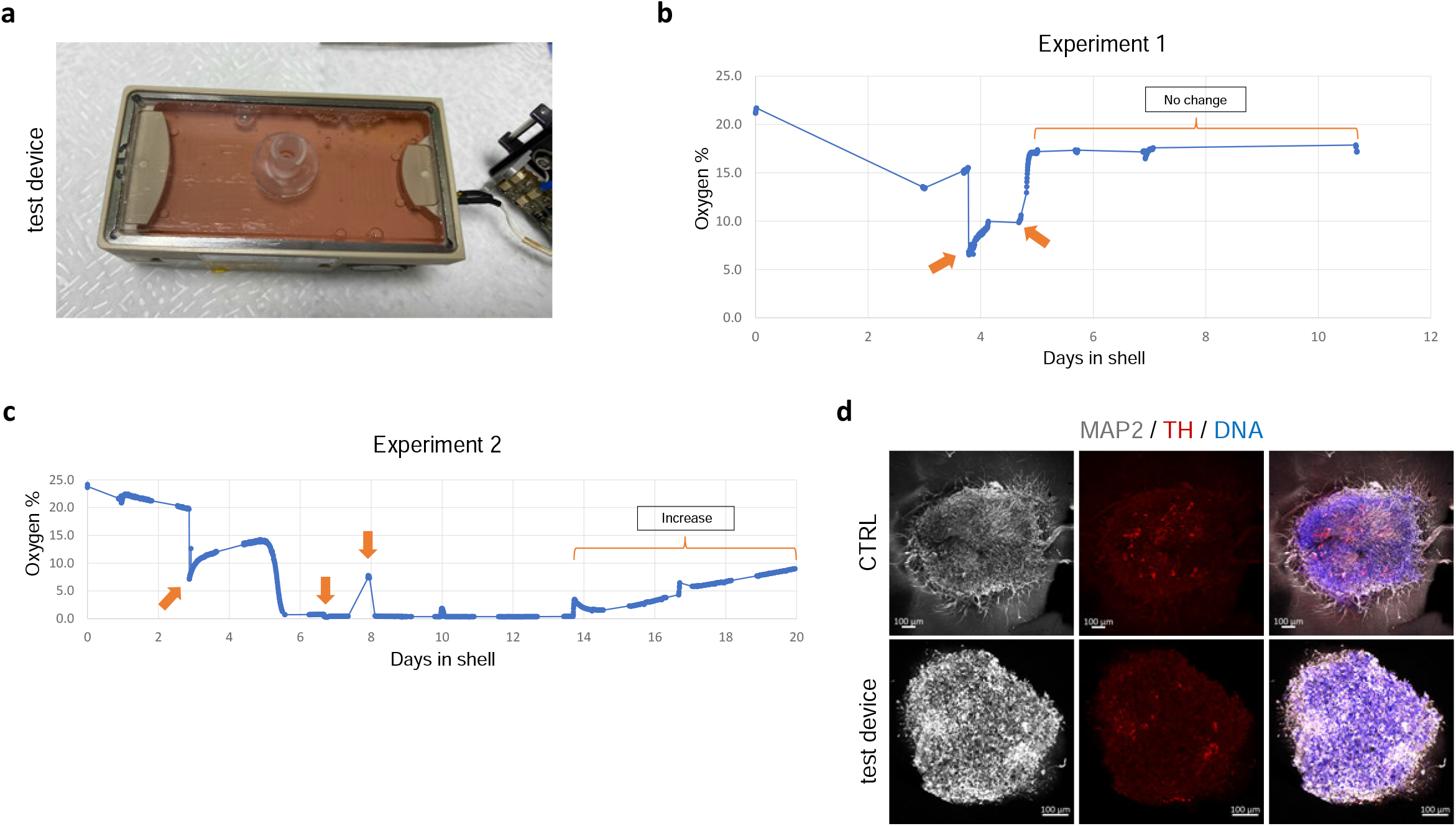
Oxygen consumption of MOs cultured in the test device. (a) Image of test device filled with MOs and culture medium. Oxygen levels were monitored via an O_2_ sensor attached to the lid of the test device. (b) Oxygen concentration within the test-device over a 10-day period. The x-axis denotes time (days 0–12), and the y-axis indicates oxygen concentration (%), ranging from 0% to 25%. Notable events are shown with the arrow: an attempted media exchange with associated leakage at approximately day 4, device reassembly and refilling at day 6, and a stable period with no changes in oxygen concentration between days 7 and 10. (c) Oxygen concentration within the test-device over a 20-day period. The x-axis represents time (days 0–20), and the y-axis shows oxygen concentration (%), ranging from 0% to 100%. Notable events are shown with the arrow: media exchange attempt with tube expansion and leakage on Day 3, a second media exchange attempt with similar observations on Day 7, and device reassembly and refilling on Day 8. A notable increase in oxygen concentration is observed between Days 14 and approximately Day 18. (d) Representative immunofluorescence images of MOs cultured for 20 days in the test device. Organoid sections were stained for TH (tyrosine hydroxylase), MAP2 (neuronal marker), and nuclei (Hoechst 33342). Scale bars: 20x, 100μm.

In conclusion, long-term MO culture in the hardware is feasible but requires optimized fluidics and oxygen control to maintain viability. These results highlight the need for careful device assembly for autonomous experiments in microgravity.

These biological test procedures were conducted to determine optimal conditions for MO survival throughout the spaceflight experiment. We evaluated control versus aged culture medium, selecting aged medium with biweekly exchanges (2x per week) for sustained viability. Fixation protocols using 4% PFA at RT preserved morphology over extended periods. Simulated hypergravity and vibration tests demonstrated that organoids withstand launch stresses without damage. Oxygen consumption analysis confirmed healthy morphology, though limited by the fixed design of the Type-IV device, indicating future media modifications may enhance oxygenation. Based on these findings, the CubeLab™ was modified to accommodate the devices for a successful MO launch to the ISS.

### Pre-launch procedures and post-flight assessment of organoid viability

Pre-launch procedures involved the shipment of both neuroepithelial stem cells (NESCs) and pre-generated midbrain organoids, together with all consumables, from the LCSB in Luxembourg to the Space Station Processing Facility (SSPF) at Kennedy Space Center (KSC) in Florida, USA. Due to the postponement of the SpaceX CRS-27 launch to the 14^th^ of March 2023, a new batch of MOs was generated at the SSPF from the shipped NESCs. Ten midbrain organoids were placed in each Yuri Type-IV device (yuri003 and yuri004) on day 8, coinciding with the switch to differentiation medium and the onset of neurite outgrowth. Afterwards, the CubeLab™ was sealed following NASA hardware air-tightening protocols. Supplementary Table 2 provides a detailed overview of pre-launch procedures, including handling steps, media exchange schedules, and the return-transport timeline after mission completion.

The CubeLab was integrated into the SpaceX Cargo Dragon capsule and powered through the Space Tango PAUL-2 locker during launch. The launch of the SpaceX Falcon 9 CRS-27 took place at launch complex 39A of the Kennedy Space Centre in Cape Canaveral, Florida, USA. The experiment took place aboard the International Space Station (ISS), and upon completion, after which the CubeLab™ was transferred back to the Cargo Dragon capsule for re-entry and splashdown. During the return phase and subsequent shipment back to Luxembourg, the CubeLab™ was maintained at 4°C, and the target post-flight temperature range of + 2°C to + 8°C was consistently preserved (Supplementary Figure 4a).

However, software errors caused major deviations from the planned autonomous operations. Only one media exchange occurred on day 3, and the software failed to reach the target temperature of 20°C for fixation. A secondary software malfunction reheated the CubeLab™ to 37°C, thereby preventing fixation (Supplementary Figure 4b-c). Consequently, the midbrain organoids returned to Luxembourg unfixed and stored at 4°C. Upon CubeLab™ opening, device yuri004 showed no leakage, and all 10 midbrain organoids were successfully recovered (Supplementary Figure 5a). In contrast, device yuri003 exhibited leakage from the cell culture chamber (Supplementary Figure 5b) and 9 midbrain organoids were retrieved. All organoids were imaged for size, fixed overnight at 4°C, and culture medium was collected for LDH assays. Due to the device leakage, widespread cell death was observed surrounding the midbrain organoids in the Geltrex matrix (Supplementary Figure 5c), accompanied by a significant reduction in size (Supplementary Figure 5d). Due to this leakage-related damage, organoids from yuri003 were excluded from further analysis.

To determine whether the observed damage reflected spaceflight-specific effects or device-related stress, we performed three ground control experiments (GE1-GE3) under identical conditions to the ISS experiment. These included one media exchange on day 3, incubation in the identical hardware for the full experimental duration, absence of external gas exchange, and the same delayed post-mission fixation. Similar to yuri003, all ground controls displayed dead cells localized within the Geltrex matrix. GE1 showed the most severe phenotype and was excluded. GE2 and GE3 maintained organoid sizes comparable to laboratory standard controls (CTRL) (Supplementary Figure 5c-d), although both exhibited significantly elevated LDH release relative to CTRL and yuri004 (ISS) (Supplementary Figure 5e), indicating increased cytotoxicity. Due to this pronounced degeneration in hardware-matched ground samples, subsequent analyses were restricted to intact flight organoids from yuri004 (ISS) and laboratory standard controls (CTRL) maintained under routine culture conditions.

### Morphological adaptations and neuronal vulnerability in ISS-midbrain organoids

Midbrain organoids cultured aboard the International Space Station (ISS) were analysed to assess growth and viability under prolonged exposure to microgravity within mission-associated constraints. Morphological and viability-related changes were evaluated following long-term spaceflight environment.

Despite the malfunction preventing scheduled media exchanges during the 40-day mission, ISS-MOs survived and exhibited robust neurite outgrowth comparable to the control MOs cultured under standard conditions (Figure 5a). Quantitative analysis of the organoid area revealed a significant reduction in size in ISS-MOs compared to controls (Figure 5b), likely reflecting cumulative metabolic constraints associated with limited media exchange. However, while nutrient-restricted conditions could have contributed to this reduction, the absence of similar effects in the ground controls, which were exposed to the same media exchange schedule but not microgravity, suggests that spaceflight-induced microgravity also likely plays a role in these changes. Despite their smaller size, LDH measurements showed no significant increase in cytotoxicity (Figure 5c), indicating that overall membrane integrity remained largely preserved under prolonged spaceflight-associated stress.

**Figure 5.**
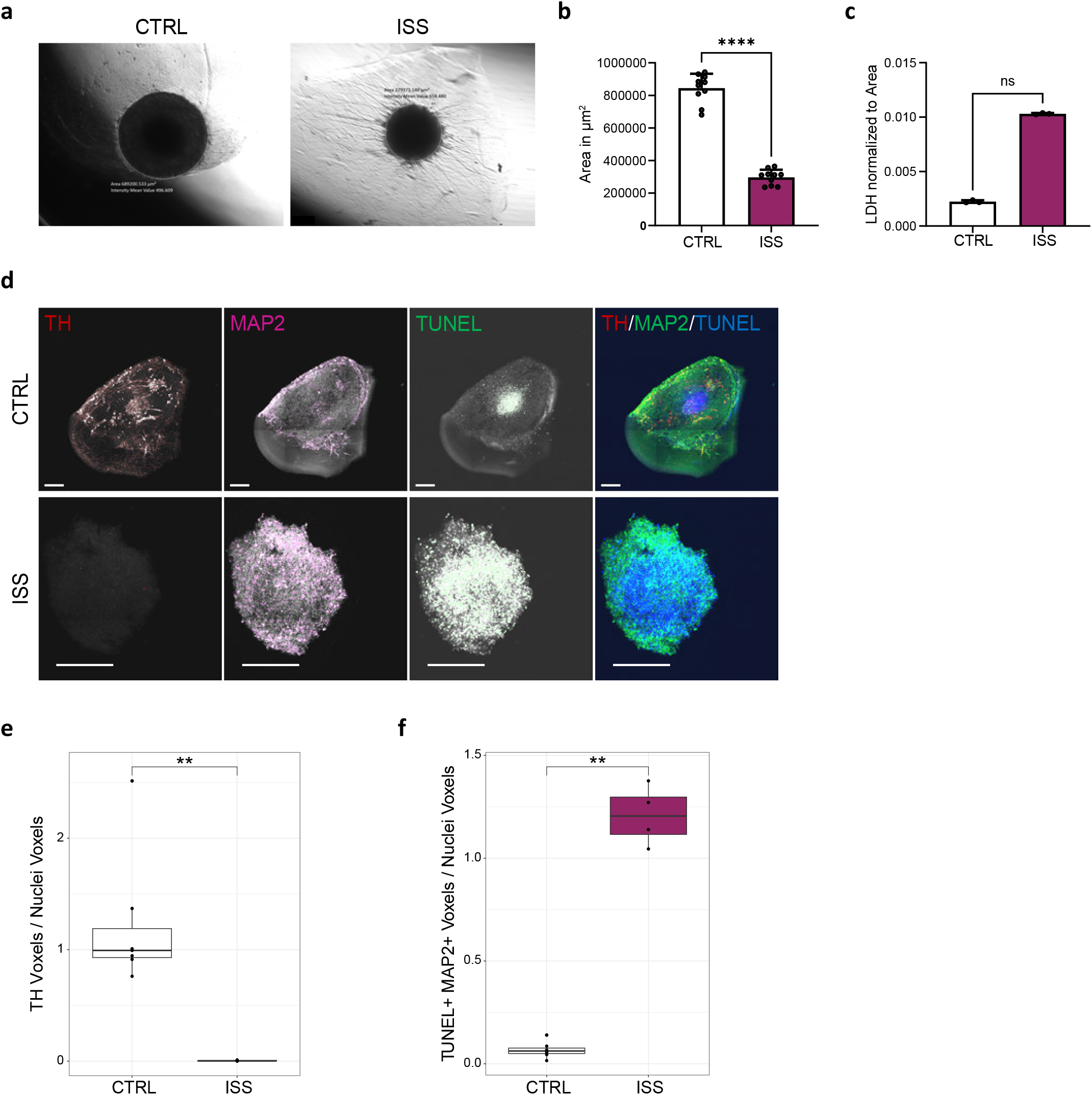
Effects of microgravity on morphology and neuronal integrity in MOs cultured on the ISS. (a) Representative brightfield images of CTRL and ISS-cultured MOs with the annotation of the Area in μm^2^ and the intensity mean value. (b) Quantification of Area in μm^2^ of 10 CTRL and 10 ISS (yuri004) MOs. Statistical significance was assessed using the Mann-Whitney *U* test. Data are mean ± SD; ****p<0.0001. (c) Quantification of cytotoxicity by LDH release assay (% normalized to control) of 10 CTRL and 10 ISS (yuri004) MOs. Statistical significance was assessed using the Mann-Whitney *U* test. Data are mean ± SD; ns = not significant. (d) Representative high-content microscope images of CTRL and ISS-cultured MO sections stained for TH (tyrosine hydroxylase), MAP2 (neuronal marker), and TUNEL (cell death marker). Grey-scaled images show masks that were created with MATLAB for quantification. Scale bars: 20x, 10011μm. (e-f) High-content automated image analysis of immunofluorescence stainings showing the foldchange of TH-positive voxels (e) and the foldchange of TUNEL-positive cells co-localizing with MAP2-positive neurons (f), normalized by total nuclei. Data are presented as boxplots. Statistical significance was assessed using the Mann-Whitney *U* test; **p < 0.01.

Immunofluorescence staining showed a notable reduction of dopaminergic neurons (TH) in ISS-MOs, whereas MAP2-positive mature neurons were comparatively preserved (Figure 5d). Quantification confirmed a significant decrease of TH signal intensity (Figure 5e), indicating selective vulnerability of dopaminergic neurons under spaceflight conditions. In contrast to the unchanged LDH levels, ISS samples showed significant increase in TUNEL-positive and MAP2-positive neurons (Figure 5f), suggesting that although overall cytotoxicity remained not significantly increased in the LDH assay, certain neuronal populations experiencing stress exhibited increased susceptibility to apoptosis.

Together, these results indicate that prolonged spaceflight exposure under nutrient-restricted conditions is associated with selective dopaminergic neuron vulnerability, while allowing partial survival and structural maintenance of other neuronal populations.

### Transcriptomic profiling reveals dysregulation of neuronal pathways in midbrain organoids cultured on the ISS

To gain deeper insight into the molecular changes associated with microgravity, we performed bulk RNA sequencing on fixed ISS and standard control (CTRL) midbrain organoids. Correlation analysis of transcriptomes revealed low similarity (r = 0.13) between ISS-MOs and CTRL (Figure 6a), indicating pronounced transcriptional divergence. Differential expression analysis identified numerous significantly up- and downregulated genes (log_2_FC ≥ 2, p.adjust < 0.05), including key regulators of cytoskeletal dynamics, vesicle trafficking, and neuronal plasticity (Figure 6b). Gene set enrichment analyses highlighted coordinated dysregulation of neuronal pathways. Reactome terms were enriched for axon guidance, nervous system development, and membrane trafficking (Figure 6c), GO analysis emphasized synapse organization, cytoplasmic translation, and neuron projection morphogenesis (Figure 6d). KEGG pathways revealed alterations in dopaminergic synapse signalling, ER protein processing, and neurodegeneration-related pathways (Figure 6e). Notably, the enrichment of dopaminergic synapse pathways is consistent with the loss of TH-positive neurons observed in the morphological analyses (Figure 5d and e). Fold change analysis of gene sets related to neuronal projection, synapse formation, and cell-death responses further supported selective neural adaptation and survival mechanisms to the ISS environment (Figure 6f).

**Figure 6.**
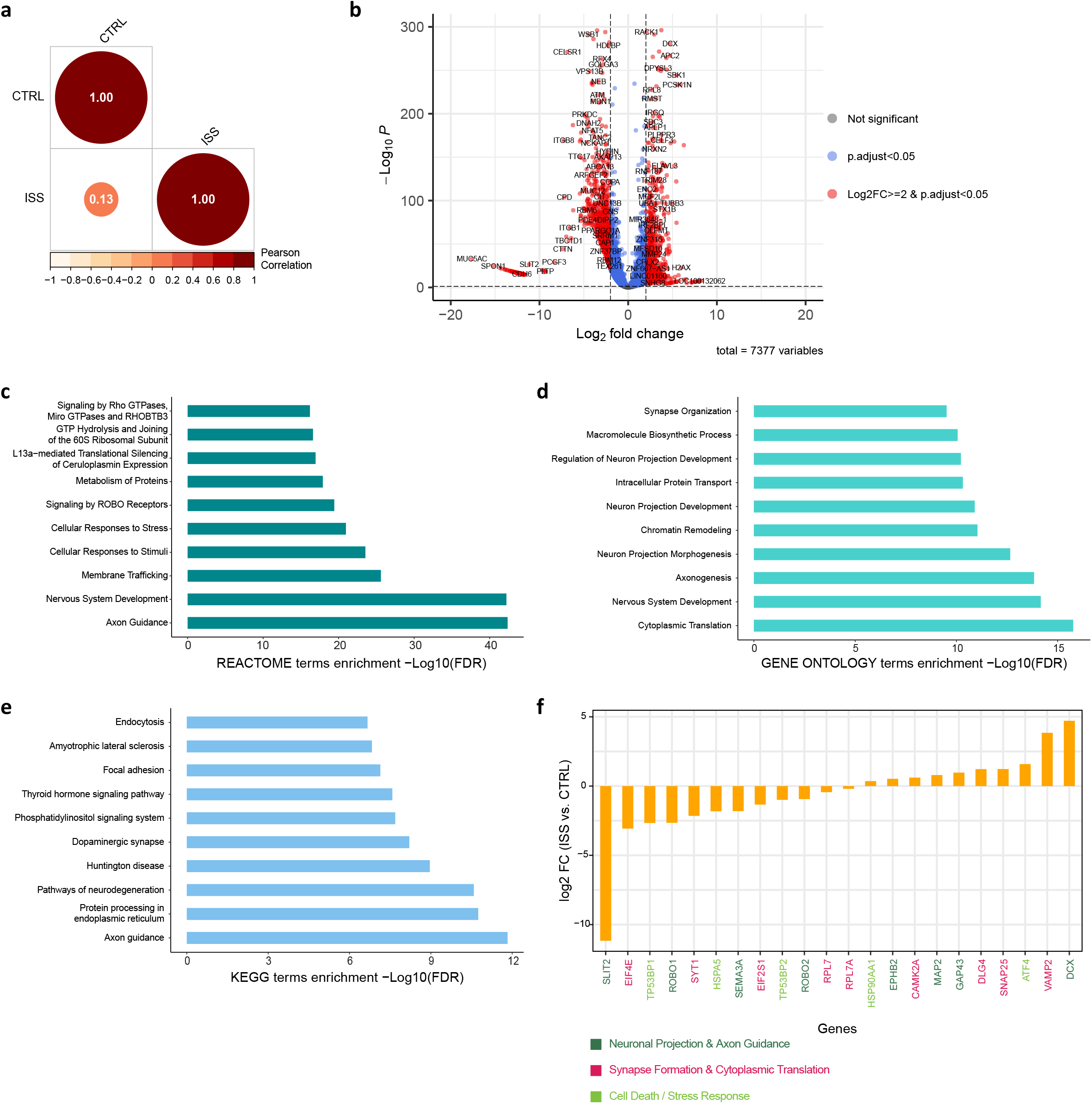
Transcriptomic profiling identifies dysregulated neuronal pathways in MOs exposed to microgravity. (a) Pearson Correlation of log2 fold changes (FC) of significant genes (p < 0.05) between CTRL and ISS MOs. (b) Volcano plot showing differential gene expression (DEGs) between ISS and CTRL MOs. Genes with log_2_FC>2 and p.adjust <0.05 are highlighted. (c) Reactome pathway enrichment analysis of the DEGs (p.adjusted <0.05) between ISS and CTRL samples, showing the top 10 significantly enriched pathways based on EnrichR results. (d) GO pathway enrichment analysis of the DEGs (p.adjusted <0.05) between ISS and CTRL samples, showing the top 10 significantly enriched pathways based on EnrichR results. (e) KEGG pathway enrichment analysis of the DEGs (p.adjusted <0.05) between ISS and CTRL samples, showing the top 10 significantly enriched pathways based on EnrichR results. (f) Log_2_ fold changes (FC) of predefined genes associated with neuronal projection and axon guidance (darkgreen), synapse formation and cytoplasmic translations (magenta), or cell death / stress response (lightgreen).

These results demonstrate that microgravity and nutrient deprivation induce a coordinated transcriptional program promoting neural plasticity, cytoskeletal remodelling, and stress resilience. The transcriptomic changes align with immunofluorescence data, including dopaminergic neuron loss, partial neuronal preservation, and maintained neurite outgrowth. ISS organoids activated survival and remodelling pathways, enabling structural maintenance despite prolonged environmental stress.

## Discussion

This study was conducted within the framework of the “Überflieger 2” student competition and therefore differs in scope and level of control from a conventional research project. While this setting provided a unique opportunity to perform a spaceflight experiment with human midbrain organoids, it also introduced inherent constraints in experimental design, robustness, and contingency handling. In particular, the combination of intended microgravity exposure and unintended hardware malfunctions represents a major limitation for data interpretation, as the observed effects cannot be unambiguously attributed to either microgravity, nutrient deprivation caused by impaired medium exchange, or the interaction of both factors. Despite these limitations, the study provides valuable insights. The experimental outcomes highlight critical aspects of process optimization, including hardware reliability, fluidic control, and environmental stability, which are essential for long-term autonomous culture in space. These findings establish an important technical and conceptual foundation that will enable more controlled and interpretable experiments in future missions. This study demonstrates the general feasibility of autonomously culturing human midbrain organoids in the Yuri Type-IV bioreactor system under both simulated and actual spaceflight conditions. Ground-based optimization established that aged medium with twice-weekly partial exchanges, together with 4% PFA fixation at room temperature, preserved organoid viability and structural integrity over extended culture. Simulated hypergravity and mechanical vibration assays further confirmed that MOs tolerated the mechanical stresses expected during launch.

Despite major hardware malfunctions during the ISS mission, including the absence of scheduled media exchanges and the failure of the automated fixation sequence, ISS-cultured organoids survived for 40 days in microgravity and maintained robust neurite outgrowth. Nonetheless, they displayed a substantial reduction in size, likely reflecting nutrient depletion and metabolic stress. Immunofluorescence analysis revealed a selective loss of TH-positive dopaminergic neurons, while MAP2-positive mature neurons remained comparatively preserved. This pattern is consistent with previous observations of dopaminergic vulnerability under metabolic and environmental stress (Popova et al., 2020; Ali et al., 2025). Transcriptomic profiling revealed extensive remodeling of neuronal pathways in ISS-MOs, including downregulation of dopaminergic synapse and neurodegeneration-related pathways, and upregulation of cytoskeletal reorganization, neuronal projection, and plasticity-associated genes. Increased apoptosis markers in neuronal subpopulations further underscored the selective cellular stress imposed by the microgravity environment. The combination of reduced organoid size, selective dopaminergic neuron loss, cytoskeletal remodelling, and increased apoptotic signatures suggests that microgravity may impose a bioenergetic and structural adaptation program in midbrain organoids. Dopaminergic neurons are characterized by high metabolic demand, extensive axonal arborization, and elevated basal oxidative stress, making them particularly vulnerable to altered nutrient diffusion, mitochondrial stress, and energy depletion (Pacelli et al., 2015; Ni & Ernst 2022). Thus, the upregulation of projection- and plasticity-associated genes in ISS-MOs likely reflects adaptive cytoskeletal and synaptic remodelling in surviving neuronal populations rather than uniform degeneration.

These results align with previous reports showing altered neuronal differentiation and cytoskeletal dynamics in microgravity (Zu Eulenburg, *et al*., 2021; Wuyts *et al*., 2025; Mattei *et al*., 2018; Marotta *et al*., 2024) and complement findings from Shaka *et al*. (2022), who demonstrated that space microgravity alters neural stem cell division, potentially influencing neuronal differentiation and survival. Our findings are consistent with prior observations of altered progenitor fate and cytoskeletal remodelling in neural organoids under microgravity (Mattei *et al*., 2018; Marotta *et al*., 2024), while extending these insights to a midbrain-specific model and revealing pronounced dopaminergic vulnerability, suggesting region- and lineage-specific sensitivity to spaceflight conditions. Together, these data suggest that microgravity-induced changes at the progenitor or early neuronal level may contribute to the dopaminergic neuron vulnerability observed in our organoids.

Importantly, our data provide a strong rationale to focus future experiments on Parkinson’s disease (PD) models. PD pathogenesis critically involves dopaminergic neuron degeneration and mitochondrial dysfunction, processes potentially exacerbated by spaceflight-induced oxidative stress and altered cellular homeostasis (Ali *et al*., 2025). Beyond PD, neurodegenerative diseases share overlapping mechanisms of metabolic stress, protein homeostasis, and cytoskeletal dysfunction, suggesting that spaceflight-based organoid models could also be extended to other disease contexts. Using disease-specific organoids will enable mechanistic exploration of how microgravity and cosmic radiation impact neurodegenerative processes, opening translational avenues relevant both to space health and terrestrial PD research.

For improved experimental outcomes, future experiments should incorporate media formulations optimized for oxygen and CO_2_ balance, such as BrainPhys, which support neuronal survival and synaptic function under challenging conditions (Saglam-Metiner *et al*., 2023). Enhanced oxygen strategies, tighter fluidic control and real-time monitoring will be essential to maintain metabolic activity for long-duration autonomous culture. To increase robustness and reproducibility, multiple biological replicates, including wild-type and disease-model organoids, should be launched in parallel using malfunction-free hardware.

Several limitations should be considered when interpreting these findings. First, the absence of scheduled media exchanges during the ISS mission precludes definitive separation of microgravity-specific effects from those arising due to nutrient depletion and metabolic stress. Second, the limited number of biological replicates restricts assessment of inter-organoid variability and reduces statistical power. Midbrain organoids generally exhibit high intra- and inter-batch reproducibility (Zuccoli *et al*., 2025), underscoring the importance of including sufficient replicates to fully leverage the stability of this model system. Third, because only one flight device (yuri004) yielded intact organoids and ground hardware-matched controls degenerated severely, transcriptomic analyses necessarily relied on standard laboratory controls rather than hardware-matched comparators, which may introduce baseline differences unrelated to microgravity. Finally, the long post-flight delay prior to fixation introduces unavoidable confound related to prost-landing cold storage. Future missions incorporating controlled fluidic exchange, multiple independent devices, and parallel ground controls subjected to equivalent nutrient distribution will be essential to disentangle these variables and more precisely isolate microgravity-driven biological effects.

In summary, despite technical limitations and hardware failures, this study provides valuable insights into how human midbrain organoids respond to long-term microgravity. These findings establish a framework for future, more controlled missions using brain organoid platforms to investigate neurodevelopmental and neurodegenerative disease mechanisms under spaceflight conditions.

## Data availability

All original and processed data as well as scripts that support the findings of this study are publicly available at this https://doi.org/10.17881/av9z-4v06. Gene expression datasets can be accessed on Gene Expression Omnibus (GEO) under the accession code GSE301053.

## Code availability

All scripts used to obtain, analyse and plot the data are available at https://gitlab.com/uniluxembourg/lcsb/developmental-and-cellular-biology/BRAINS_2025.

## Acknowledgements

This work was mainly supported by the Überflieger 2 program organized by the German Space Centre (DLR), Luxembourg Space Agency (LSA), Deutsche Physikalische Gesellschaft (DGP) and Yuri GmbH. We thank the Developmental and Cellular Biology Group at the Luxembourg Centre for Systems Biomedicine (LCSB) for providing the consumables for the biological part, as well as their valuable input. We acknowledge the Space Robotics Group at the (SnT) for their valuable contributions, and OrganoTherapeutics (OT) for sharing the wild-type (WT) cell line with us. Further, we acknowledge the sponsoring support from our bronze sponsor, Foyer Assurances S.A. Luxembourg and our travel sponsors, Banque Internationale à Luxembourg (BIL), Technoport S.A. Luxembourg, Amis de l’Université de Luxembourg and the Société Européenne des Satellites (SES). We thank Dennis Balasus from Yuri GmbH for the help during the preparation of the mission in Luxembourg and in the Space Station Processing Facility (SSPF) on the NASA site, Elizabeth S. Carter from NASA for onsite help and Dr. Devin Mair, from the Johns Hopkins University and stationed at the SSPF for the generous donation of last-minute consumables for our project. We would also like to thank Bob Lamboray (LSA) for supporting the project since the first proposal and the other Überflieger 2 participants of the ADDONNIS, Glücksklee and FARGO groups. We would like to thank the LCSB Bioimaging Platform for supporting high-content imaging and image analysis workflow. This work leading to this manuscript was supported by the following funding: i2TRON Doctoral Training Unit and the Fonds National de la Recherche (FNR) Luxembourg i2TRON PRIDE19/14254520. For the purpose of Open Access, the author has applied a CC BY public copyright license to any Author Accepted Manuscript (AAM) version arising from this submission.

## Author contributions

All authors contributed to the review and search of the literature. Conceptualisation, E.Z., D.V.G., A.C.C., L.M.A., J.C.S., and M.O.M., Investigation, E.Z., D.V.G., A.C.C., L.M.A., and J.D.C., Writing and original draft preparation, E.Z., Review and editing, E.Z. and J.C.S. Visualisation, E.Z., D.V.G., A.C.C., L.M.A., and J.D.C, Project administration, E.Z., D.V.G, J.C.S., and M.O.M, Funding acquisition, E.Z., D.V.G., A.C.C., L.M.A., J.D.C., J.C.S., and M.O.M. All authors have read and agreed to the published version of the manuscript.

## Competing interests

J.C.S. declares no competing non-financial interests but declares competing financial interests as cofounder and shareholder of OrganoTherapeutics société à responsabilité limitée (SARL). The remaining authors declare no competing interests.

## Figure legends

**Supplementary Figure 1. Pre-flight validation of electronic subsystems**. (a) Test circuit 1 validating temperature-sensor readings using a thermistor connected to an Arduino Micro (Rev3). (b) Test circuit 2 verifying heater control using an NPN transistor (s8050) driving a MOSFET (IRFZ44) to switch the 12V heating element. (c) Test circuit 3 assessing bidirectional motor control using L298N H-bridge driver connected to the Arduino.

**Supplementary Figure 2. Effect of media freshness and exchange frequency on MO viability and differentiation**. (a-c) Representative immunofluorescence images of MOs at day 10 (a), day 20 (b) and day 30 (c) cultured under different conditions: control medium (freshly prepared at 4°C) with standard exchange (CTRL), or with partial exchange twice (2x) or once (1x) per week, and aged medium (stored at 37°C) with the same exchange frequencies. Organoid sections were stained for TH (tyrosine hydroxylase), MAP2 (neuronal marker), and nuclei (Hoechst 33342). Scale bars: 20x, 100 μm. (d) Quantification of cytotoxicity by LDH release assay (% relative to positive control) shows significantly elevated cell death in aged media and reduced exchange conditions across all time points. Statistical significance was assessed using an ordinary two-way ANOVA with Dunnett’s multiple comparison test. Data are mean ± SD; ***p < 0.001, ****p < 0.0001; ns = not significant.

**Supplementary Figure 3. Optimization of fixation conditions for MOs under simulated spaceflight constraints**. (a–b) Representative immunofluorescence images of MOs collected at day 20 (a) and day 30 (b), fixed with 1%, 2%, or 4% paraformaldehyde (PFA) for 40 or 50 days at 4°C. Organoid sections were stained for TH (tyrosine hydroxylase), MAP2 (neuronal marker), and nuclei (Hoechst 33342). (c–d) MOs fixed ON (overnight) at either 4°C or room temperature (RT) at day 10 (c) or day 17 (d) to simulate pre-fixation conditions aboard the ISS. Scale bars: 20x, 100μm.

**Supplementary Figure 4. Environmental monitoring and software performance of MO experiment during transport and ISS culture**. (a) Temperature results of average temperature of the package during transport from the US to Luxembourg. Results show the monitor configuration, the recorded data, the monitor read and the summary data. (b) Timeline of average temperature of the devices during the ISS experiment. Shell 1 corresponds to device yuri004 and shell 2 to device yuri003. The data was stored on an SD card during the experiment, that was retrieved upon arrival and evaluated. (c) Overview of software errors that occurred during the ISS experiment, highlighted in red-outlined boxes. Three timers were implemented; the errors occurred within the nutrient exchange and the fixation code.

**Supplementary Figure 5. Evaluation of device integrity, organoid morphology, and cytotoxicity following ISS experiment**. (a) Image of yuri004 device post ISS-experiment. The device showed no leakage and 10 MOs were retrieved from the device. (b) Image of yuri003 device post ISS-experiment. The device leaked from the cell culture chamber which had a huge bubble. (c) Representative brightfield images of CTRL, ISS-cultured MOs from devices yuri004 and yuri003 as well as the three ground control experiments (GE1-GE3) with the annotation of the Area in μm^2^ and the intensity mean value. (d) Quantification of Area in μm^2^ of 10 CTRL, 10 ISS_yuri004, 10 ISS_yuri003 and 10 MOs of each GEs. Statistical significance was assessed using the Kruskal-Wallis with Dunn’s multiple comparison tests. Data are mean ± SD; **p < 0.01, ***p < 0.001, ****p<0.0001. (e) Quantification of cytotoxicity by LDH release assay (% normalized to control) of 10 CTRL and 10 ISS (yuri004) MOs. Statistical significance was assessed using the Kruskal-Wallis with Dunn’s multiple comparison tests. Data are mean ± SD; **p < 0.01, ns = not significant.

