## Supplementary Figures for "Microgravity enhances the viability of midbrain organoids on the International Space Station"

Supplementary Figure 1

a

##### Test circuit 1: Thermistor test

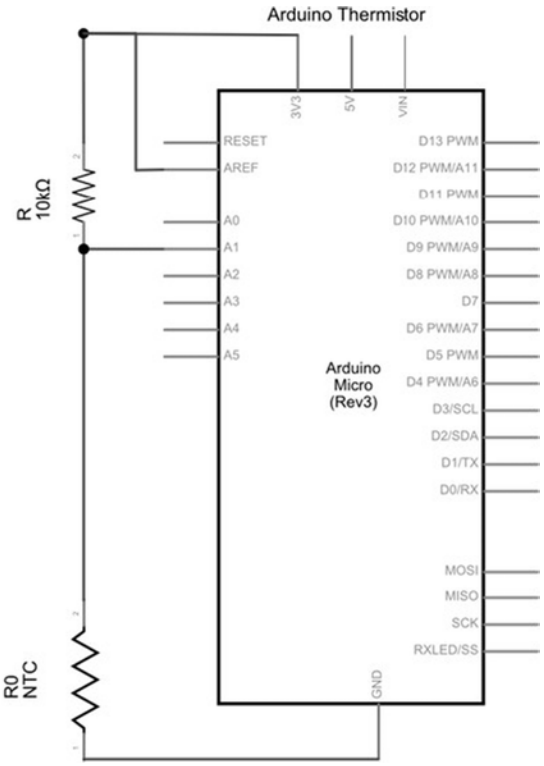


b

##### Test circuit 2: Heater test

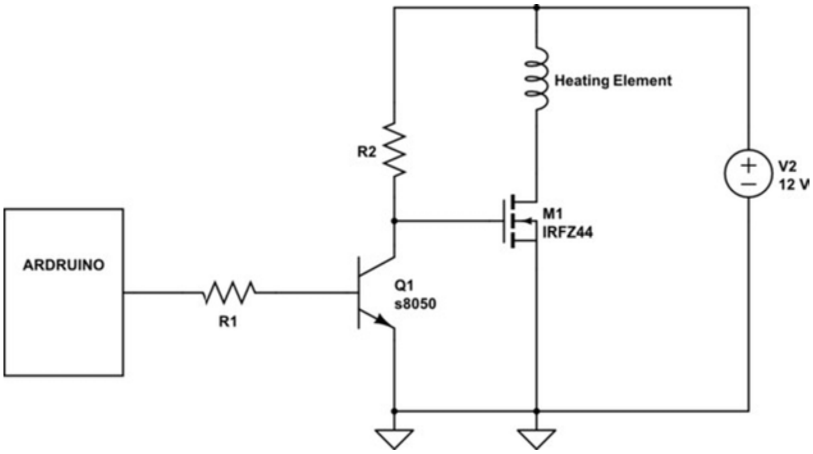


c

##### Test circuit 3: Motor test

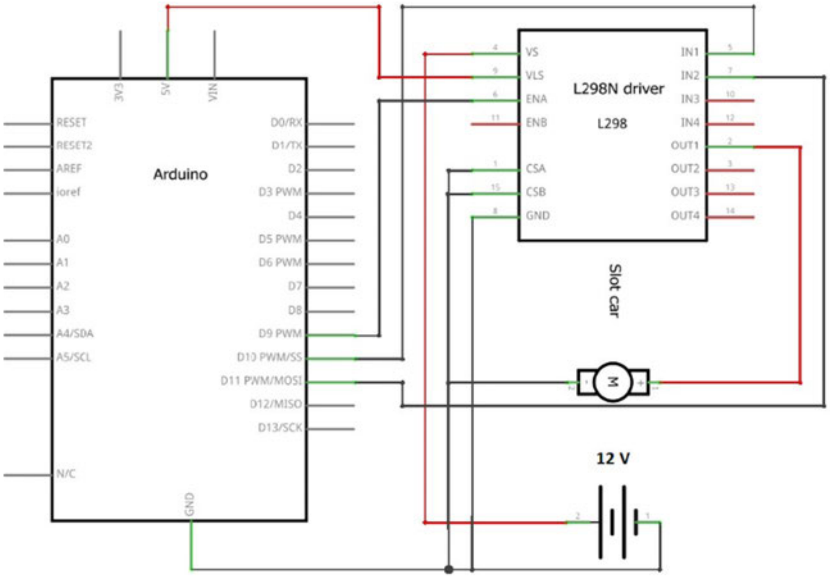

### Supplementary Figure 2

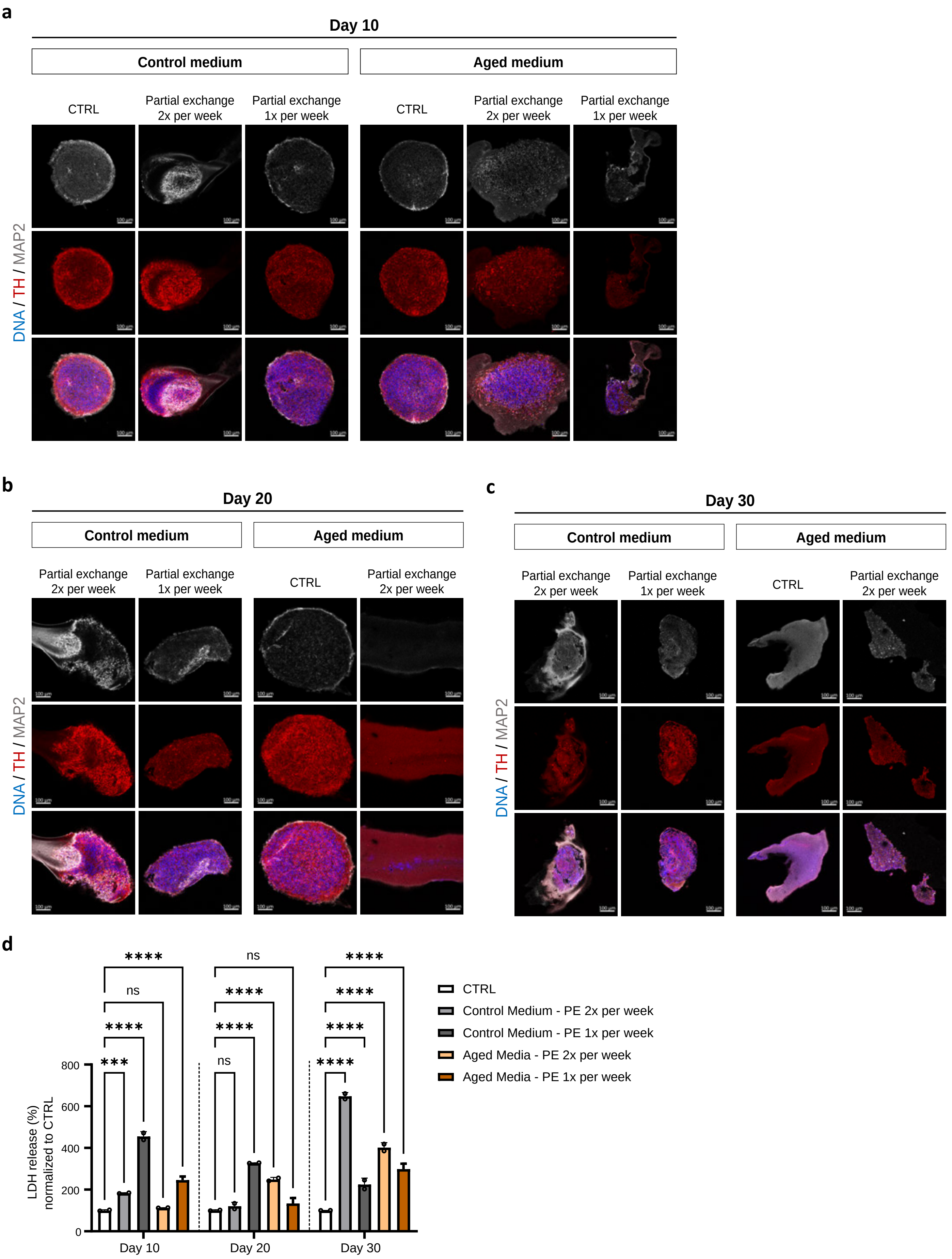

**Supplementary Figure 3**

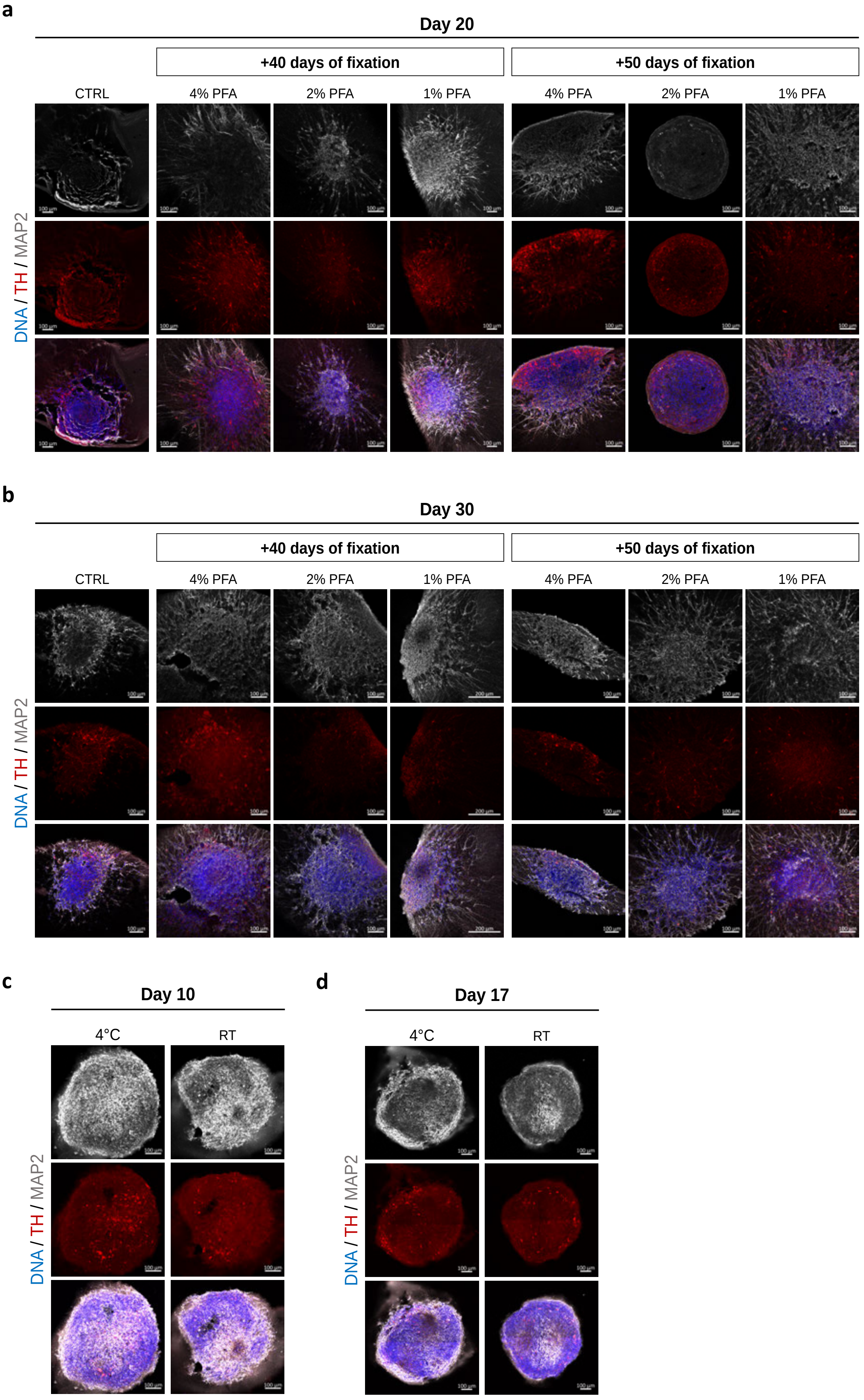

### Supplementary Figure 4

**a**

**TempTale ® Ultra**

**Serial #: KKD6N033M0**

**Monitor Configuration**

| | |
|---|---|
| Start Up Delay: | 30 Minutes |
| Interval: | 5 Minutes |

**Monitor Read**

| | |
|---|---|
| On: | 26-Apr-2023 07:40:21 AM |
| By: | A081958~worldcourier |
| Note: | All Times in GMT |

**Recorded Data**

| | |
|---|---|
| First Point: | 18-Apr-2023 13:15:56 PM |
| Stop Time: | 25-Apr-2023 07:30:51 AM |
| Number Of Points: | 1947 |
| Trip length: | 6 Days, 18 Hours, 14 Minutes, 55 Seconds |

**Summary Data**

| | |
|---|---|
| Low Extreme: | 2.50 °C @ 18-Apr-2023 01:45:56 PM |
| High Extreme: | 7.83 °C @ 18-Apr-2023 05:25:56 PM |
| Mean and Std Dev: | 5.08 ± 0.45 °C |
| Mean Kinetic Temp: | 5.09 °C |
| Activation Energy: | 83.14 kJ/mol |

**b**

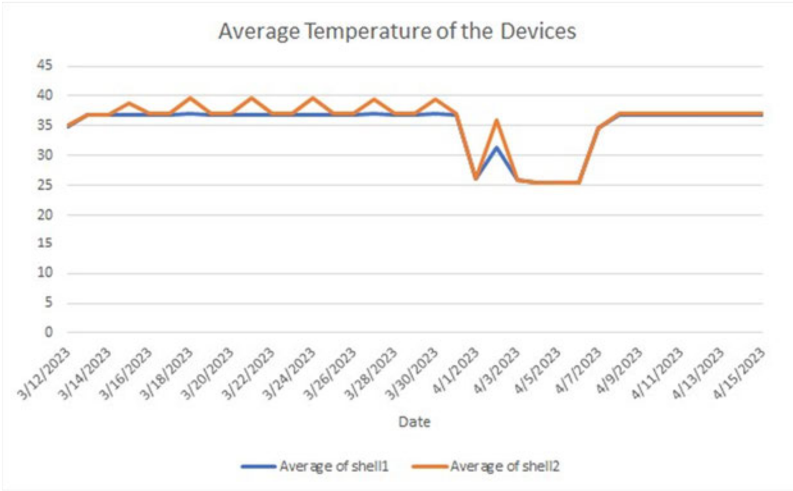


**c**

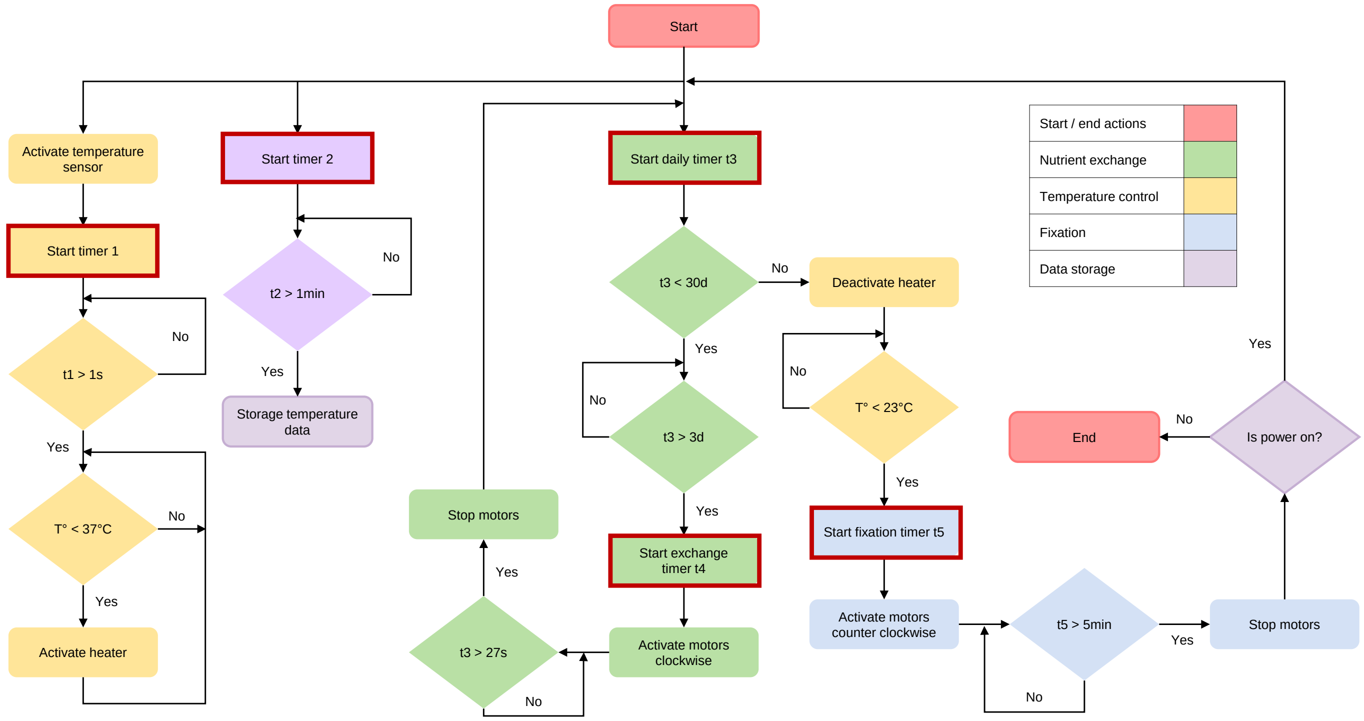

#### Supplementary Figure 5

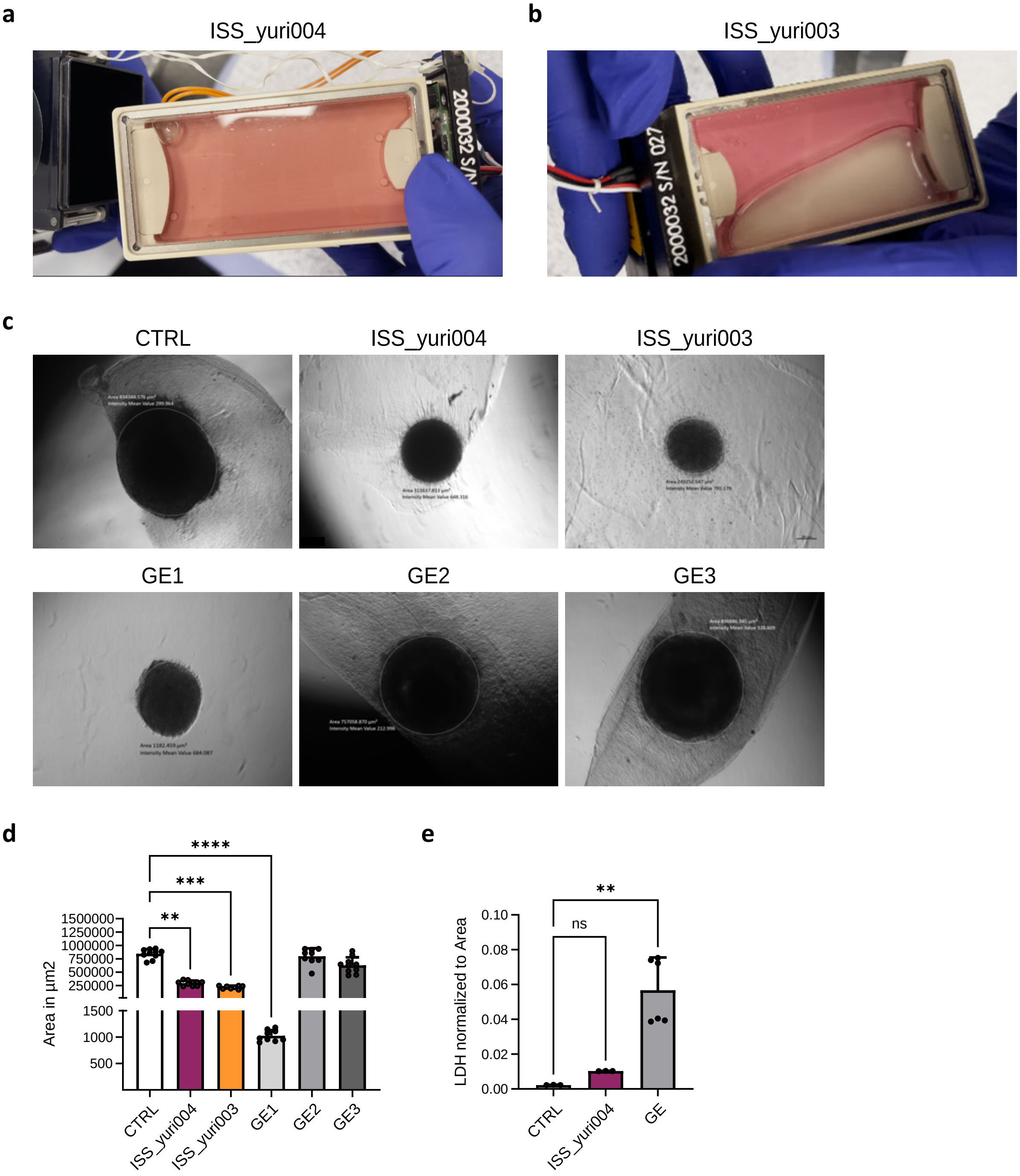
