## Supplementary Table 1 for "Microgravity enhances the viability of midbrain organoids on the International Space Station"

| Lab Identifier | Diagnosis | Patient | Sex | Age of sampling | Source of iPSC |
| --- | --- | --- | --- | --- | --- |
| 232 | Healthy | T12.9/C1-2 | F | 53 | Reinhardt et al., 2013 |

**Supplementary Table 1. Cell line used in this study**
